# Lipid A double bond position determination using ozone and laser-induced dissociation

**DOI:** 10.1101/2024.02.15.580351

**Authors:** Abanoub Mikhael, Helena Pětrošová, Derek Smith, Robert K. Ernst, David R. Goodlett

## Abstract

Ozone coupled with tandem MS is a traditional tool to assign double bond position in lipids. However, few studies were reported on the use of ozone for double bond determination of bacterial lipids. Most MS-based double bond localization methods used in bacterial lipidomics focused on low molecular weight phospholipids, but never tested for assigning double bond position in complex glycolipids such as Lipid A. Here, we present a simple approach to identify unsaturated lipid A double bond position using on-MALDI plate ozone treatment of intact Lipid A.

## Introduction

Lipid A is one of the Gram-negative bacterial virulence factors contributing to pathogenicity in mammals and even the plant kingdom. Lipid A can bind to TLR-4 receptor and initiate immune response.^1^ The binding of Lipid A to TLR-4 greatly depends on the number and the type of its fatty acyl chains.^1^ Conventionally, double bond positions of Lipid A fatty acid chains have been determined mainly using gas chromatography-mass spectrometry (GC-MS) that result in loss of information regarding the position of each fatty acid present prior to hydrolysis.^2^ The assignment of double bonds using GC-MS requires a series of chemical reactions: complete hydrolysis of lipid A to its fatty acid components, conversion of fatty acids to methyl ester, and derivatization with complex reagents such as 4,4-dimethyloxazoline (DMOX), pyrrolidides and dimethyl disulfide (DMDS). ^2^

Here, we show that ozone reacts selectively with the lipid A double bond without forming any side products. The analysis of the resulting Lipid A ozonide derivatives by MALDI induces its decomposition to products ions diagnostic to the Lipid A double bond location. Specifically, with this described method, doubles bonds can be located in the fatty acids of lipid A molecules without the loss of position concomitant with hydrolysis required for GC-MS analysis.

## Materials and Methods

The ozone treatment was performed on two synthetic mono-unsaturated lipid A (BECC 438 and 470) with previously known proposed structures ^3^, and further confirmed by highly accurate MS (∼ 1 ppm) and MS/MS (∼ 5 ppm) measurement on Q Exactive HF orbitrap equipped with Spectroglyph MALDI source (Figure 1 and 2). BECC 438 is a di-phosphorylated lipid A (DPLA), while BECC 470 is a mono-phosphorylated lipid A (MPLA).^3^ For the sake of brevity, here the ozone reaction is demonstrated on the penta-acylated form of BECC 438 and the tetra-acylated form of BECC 470. The ozone treatment was performed using the same apparatus setup previously used by **Bednařík *et al***.^4^ for on-tissue ozonation prior to MALDI-imaging. The Lipid A standards were analyzed before and after the ozone treatment using norharmane as a matrix. The analysis was performed in the negative ion mode on a timsTOF *flex* MALDI-2 instrument (Bruker, Bremen, Germany).

**Figure 1:**
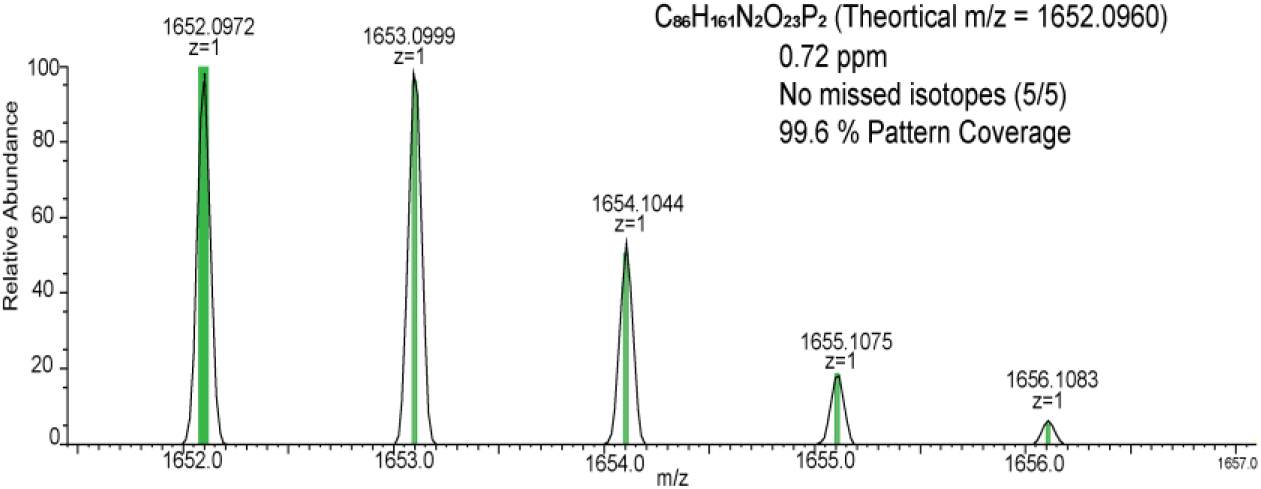
MALDI-QE accurate MS measurement of the penta-acylated form of BECC-438 (0.72 ppm and 99.6% isotopic distribution coverage)

**Figure 2:**
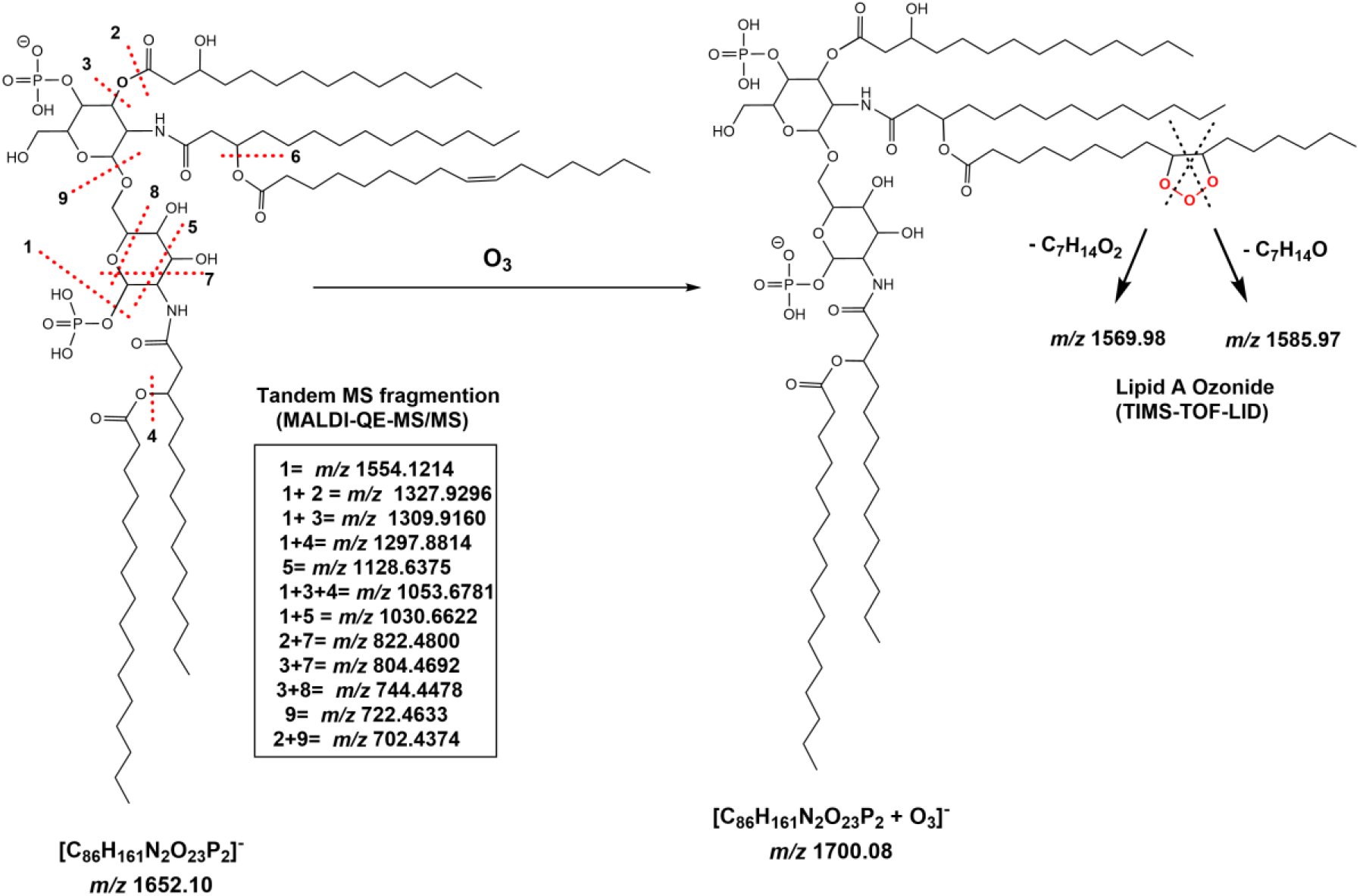
Penta-acylated form of BECC 438-DPLA pre- (left) and post- (right) ozonation. The original structure of BECC 438 (left) was confirmed with accurate MS/MS measurement on MALDI-QE as judged by the fragment ions marked with dashed lines (see inset table; tandem mass spectrum not shown), where all fragment ions were within 5 ppm. After ozonation of BECC 438, laser induced dissociation (LID) in the source of a MALDI-TIMS-TOF produced fragment ions (*m/z* 1569 and *m/z* 1585) indicative of a C16:1 double bond at position C-9.

## Results and Discussion

The penta-acylated form of the BECC 438-DPLA (*m/z* 1652.09, Figure 3-A) contains five fatty acid chains (3 x C12:0, C16:0 and C16:1). The ozone gas (O3, 47.98 Da) reacted with the single double bond in the Lipid A C16:1 chain forming the corresponding ozonide at *m/z* 1700.07 (1652.09 + 47.98). Subsequently, the MALDI laser induced the ozonide ring decomposition and formation of diagnostic products ions at *m/z* 1585.97 and 1569.98 indicating that the double bond was located at C-9 (Figure 2 and 3-B) as expected. The same strategy was also applied to the tetra-acylated form of the BECC 470-MPLA (*m/z* 1345.94, 3-C) leading to the formation of its expected corresponding ozonide (*m/z* 1393.92, Figure 3-D) followed by its laser induced decomposition to *m/z* 1263.82 and *m/z* 1279.81.

**Figure 3:**
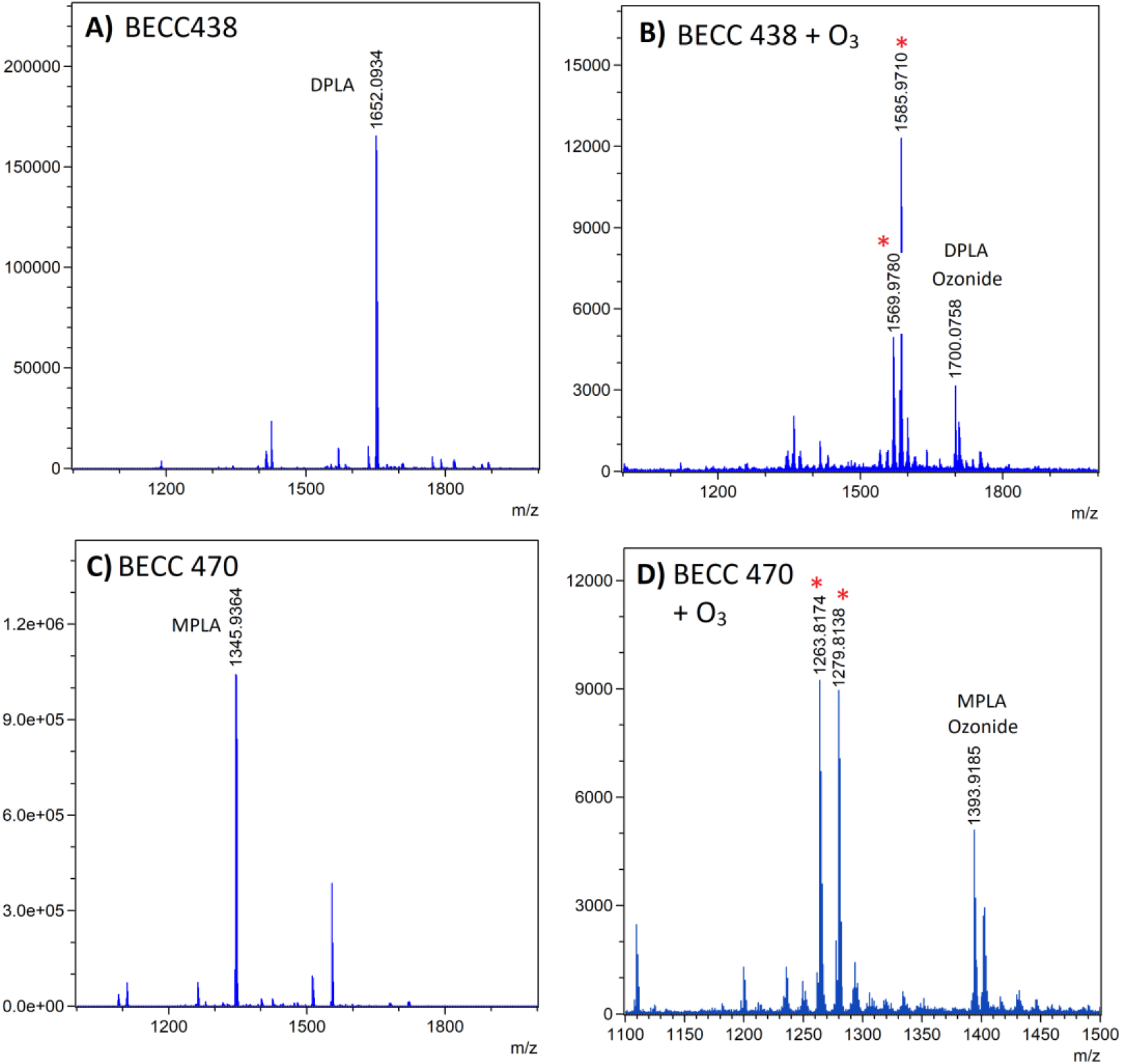
MALDI-TOF-MS of **A)** BECC438-DPLA. **B)** BECC438-DPLA post ozone treatment. **C)** BECC470-MPLA. **D)** MPLA form of BECC470 post ozone treatment. Diagnostic ions resulted from the laser-induced ozonide ring decomposition are indicated with red asterisk.

Our future work will focus on utilizing the ozone treatment to analyze different mono- and poly-unsaturated Lipid A in complex mixtures and from different bacterial sources. Defining unsaturation positions in Lipid A opens a way for new structure-activity relationships studies in the future and understanding how the number and positions of double bonds affects the Lipid A binding affinity to TLR4 and bacterial pathogenicity.

## Notes

### Competing Interest Statement

The authors have declared no competing interest.

